# Facultative social exploitation among *Myxoccocus* bacteria due to non-responsive local competition

**DOI:** 10.1101/833517

**Authors:** jeff smith, Gregory J. Velicer

**Affiliations:** No affiliation, St. Louis, Missouri, USA; Institute for Integrative Biology, ETH Zürich, Zürich, Switzerland.

**Keywords:** adaptation, cheating, competition, cooperation, *Myxococcus xanthus*, social evolution

## Abstract

Microbes cooperate in many ways, but it is unclear to what extent they also have adaptations to detect and exploit unrelated social partners. Experimental data often cannot discriminate between social exploitation caused by complex adaptive traits and exploitation caused by simple deleterious mutations. Here we demonstrate facultative social exploitation among *Myxococcus* bacteria due to simple non-responsive local competition. We show that the time dynamics of developmental exploitation by a tan phase variant are consistent with a model where cells do not respond to the presence of other genotypes but simply compete for some shared resource necessary to produce spores. The model also predicts the frequency-dependent responses of strains to genetic chimerism in multicellular fruiting bodies. Interactions between naturally occuring soil isolates, on the other hand, are consistent with strong interference competition where the winner almost completely prevents the loser from producing any spores. These results show how facultative social exploitation does not require elaborate mechanisms to detect and respond to foreign genotypes but can instead be caused by simple competition acting on a local scale. *Myxococcus* cells, like social insects, cooperate in some ways while simultaneously competing in others.

## Introduction

When researchers first began examining microbial life through the lens of social evolution, one of the earliest and strongest predictions was that microbes would have adaptations to facultatively exploit their social partners [1]. In its most sophisticated form, microbes would directly detect the presence of non-clonemates and then respond by contributing less to costly cooperation or invest more in competitive traits. These adaptations would allow microbes to benefit from cooperation when among clonemates and benefit from exploitation when among genetically dissimilar individuals—the best of both worlds.

Identifying such adaptations, if they exist, is important for at least two reasons. First, it helps us understand microbial structure and function. Nadell and colleagues [2] pointed out that while social evolution research has benefited greatly from microbiological experiments, microbiology has not equally benefited from social evolution. Using social evolution theory to identify functional traits so far overlooked by molecular microbiologists would help alleviate this disparity. Second, adaptations by definition increase organismal fitness. Identifying whether and how traits are adaptive would help us understand how those traits contribute to microbial survival and proliferation. Practically, this could help us better aid beneficial microbes or inhibit unwanted microbes. Understanding how traits contribute to microbial fitness can also help guide antimicrobial intervention strategies [3].

It is currently unclear to what extent microbes do indeed possess adaptations to responsively exploit foreign genotypes. Much of the relevant data comes from “mix experiments” that compare the fitness of strains on their own to their fitness in the presence of a competitor. In many cases, two strains will perform similarly when separate, but when mixed together one strain’s fitness will increase and the other’s will decrease [1, 4, 5, 6, 7, 8]. Some microbes also have systems that limit cell-to-cell contact between genetically distinct strains [6, 9, 10, 11], perhaps to avoid being exploited. Mixed-genotype groups sometimes perform worse than comparable single-genotype groups, possibly due to the costs of exploitation [5, 6, 12, 13]. Perhaps the clearest evidence comes from work showing that natural variation in bacterial quorum sensing causes strains to cooperate most when locally common and least when rare [14]. As suggestive as these findings may be, establishing whether and how some trait is an adaptation is notoriously difficult. As famously pointed out by George Williams and others [15, 16], biological traits often have incidental side-effects that are not their primary function—life history trade-offs that increase reproduction but decrease lifespan, for example.

Previously, we identified and theoretically developed an important alternative hypothesis to adaptive exploitation responses [17]. We showed that when there is competition within social groups for some locally shared resource, then any difference in competitiveness for that resource would have the same effect in mix experiments as responsive exploitation: mixing would increase one strain’s absolute fitness and decrease the other’s. In this scenario, cells do not react to the presence of other strains nor manipulate their behavior—it is simply the normal dynamics of competition and “soft” selection expressed within a social group. Social exploitation would thus be facultative in terms of fitness effects but not in terms of behavior. These interactions could even be caused by purely deleterious mutations that accumulate when microbes mostly interact with clonemates, relaxing selection for within-group competitiveness. Calculations using published data predicted these kinds of deleterious mutations to be common in natural populations [17].

Here we test our hypothesis using *Myxococcus* bacteria [18]. *Myxococcus xanthus* cells live in soil, where they secrete secondary metabolites that lyse other bacteria which they then consume. When starved for amino acids, *M. xanthus* cells exchange chemical signals, aggregate, and develop into multicellular fruiting bodies. Most cells in a fruiting body die, but a minority survive as stress-resistant spores. Codevelopment of naturally occuring soil isolates often increases the absolute sporulation success of one strain while decreasing the other’s [5, 6], in part due to contact-dependent interference competition [19]. Facultative exploitation also occurs among strains evolved separately in the laboratory but is not associated with faster morphological development [8].

In some cases, social exploitation by *M. xanthus* appears linked to nonproduction of yellow pigments [20]. Wild-type cells stochastically switch between the common yellow form and a rarer tan variant that becomes over-represented among spores when the two types develop together [21, 22, 23, 24]. Here we isolate a tan phase variant, show that it exhibits facultative social exploitation, and experimentally test whether that exploitation is caused by constitutive local competition for some resource necessary to produce spores. We test three predictions: (1) total group productivity of mixed-genotype groups is roughly constant with no antagonism between strains, (2) the winner is the strain that sporulates faster and/or earlier, and (3) sporulation rate and lag time are unaffected by the presence of competitors, with the main effect of codevelopment being to halt spore production when total group productivity is exhausted. We also formulate a mathematical model of local resource competition and test whether it accurately predicts how sporulation success depends on genotype frequency.

## Methods

### Strains and culture conditions

GJV1 is a wild-type derivative of laboratory *M. xanthus* strain DK1622 [25]. GJV2 is a spontaneous rifampicin-resistant mutant of GJV1 [26]. GJV10 is a kanamycin-resistant derivative of GJV1, transformed with integrated plasmid pDW79 [27]. GJV10-T is a spontaneous, swarming-proficient tan phase variant of GJV10. A23 and A47 are natural isolates from soil in Tübingen, Germany [6]. We chose these isolates because they had previously been found to produce similar numbers of spores on their own yet show exploitative interactions when developing together [6]. A23 and A47 were each previously marked with resistance to rifampicin (Rif^R^) by selecting for spontaneous mutants and separately marked with resistance to kanamycin (Km^R^) by transforming with integrated plasmid pREG1727 [6]. All strains were stored in 20% glycerol at −80 °C.

We grew cells using CTT media [28], either in liquid culture with shaking at 300 rpm or on agar plates. Where indicated, we supplemented CTT media with kanamycin at 40 µg/ml or rifampicin at 5 µg/ml. For both vegetative growth and sporulation we incubated cells at 32 °C. We incubated all plates in the dark at 90% relative humidity.

### Sporulation assay

We assayed sporulation by placing exponentially growing cells onto non-nutrient agar, where starvation induces development into multicellular fruiting bodies. To obtain cells with which to start experiments, we first inoculated strains from frozen stocks onto CTT hard agar (1.5% w/v). We then looped cells grown on these plates into CTT liquid and incubated them until they were in exponential growth (log phase). For GJV10-T, to insure the largest fraction of tan phase variants possible, we instead plated frozen plated stocks for single colonies in CTT soft agar (0.5%) and then picked a tan colony into liquid media. We centrifuged log-phase cultures at 4500 ×g for 15 minutes and resuspended cells in TPM buffer [29] to 5 × 10^9^ cells/ml. To begin the assay, we mixed resuspended cells at strain ratios 0:1, 1:99, 1:9, 1:1, 9:1, 99:1, or 1:0, then spotted 100 µl (5 × 10^8^ cells) onto TPM agar (1.5%). We incubated these plates up to 5 days, harvested cells with a sterile scalpel into 1.0 ml distilled H_2_O, and heated them for 2 hrs at 50 °C to kill any remaining vegetative cells. We dispersed spores by sonication, serially diluted them in dH_2_O, and then plated them for single colonies in CTT soft agar (0.5%). To obtain separate counts for codeveloping strains, we plated cells in CTT supplemented with kanamycin, with rifampicin, and with no antibiotic.

For experiments testing the effect of strain frequency on sporulation success, mixing frequencies were tested once per experimental block. Each of the 4–7 experimental blocks were biological replicates, performed on different days with strains independently grown from frozen stocks. The marker control competition (GJV10 + GJV2) was included in all experimental blocks.

For experiments examining the time dynamics of sporulation, each time point was a separate, destructively sampled plate. Time points across the whole course of development were taken for each of 3–4 experimental blocks that were biological replicates, performed on different days with strains independently grown from frozen stocks. The marker control competition (GJV10 + GJV2) was included in all experimental blocks except those for natural isolates (A47 + A23).

### Data analysis

We analyzed all data using the R environment for statistical computing [30]. We calculated sporulation success as the number of viable spores determined by colony counts divided by the initial number of starving cells: *w* = *n*′*/n*. This measure is equivalent to absolute Wrightian fitness over the developmental period. We tested differences in sporulation success among strains when developing on their own using Welch two-samples *t*-tests with unequal variance.

We calculated a strain’s within-group competitiveness as the ratio of its sporulation success to its competitor’s (*w*_A_*/w*_B_). We analyzed relative fitness data using multiple linear regression, fitting models to log_10_(*w*_A_*/w*_B_). When data showed heteroscedasticity, we used weighted least-squares regression as implemented in the nlme package, fitting models by maximum likelihood with an exponential relationship between variance and mixing frequency. We tested statistical significance by comparing models fit with and without the terms of interest using *F*-tests for linear models and likelihood ratio (LR) tests for weighted least-squares models. We analyzed the effect of mixing frequency on the sporulation success of both the total group and individual strains using multiple linear and weighted least-squares regression, fitting models to log_10_ *w*. For some responses we fit power functions using nonlinear least squares and nonlinear weighted least squares modelling.

We analyzed developmental time series data using the approach described by Bolker [31], fitting curves with biological meaningful parameters and making statistical inferences about those parameters. We had no prior expectation for the functional form of developmental dynamics but had good success fitting three-parameter Gompertz functions of the form

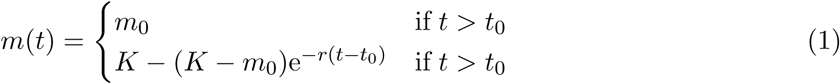

where *m*(*t*) is log_10_(spores/cell) at time *t* (hrs after starvation), *m*_0_ is a lower detection limit of log_10_(10 spores/2.5 × 10^8^ cells), *t*_0_ is the lag time until spores first become detectable, *r* is a sporulation rate parameter (/hr), and *K* is the asymptotic maximum of *m* as time goes on. The three free parameters are *r, K*, and *t*_0_.

To fit curves to data, we set data points with zero spore counts to the lower detection limit, then fit Eqn. 1 by maximum likelihood using the `mle2` command from the *bbmle* package in R. We used quasi-Newton optimization with normally distributed *m*(*t*) error and standard deviation *σ* (another free parameter). We determined likelihood profiles to check for the presence of nearby alternative minima. To determine how a strain’s developmental dynamics were affected by mixing, we fit models to data for that strain both separate and in codevelopment, then tested the significance of a mixing factor on *r* and *t*_0_. To compare strains, we fit models to both their data simultaneously and then tested the significance of a strain factor on *r* and *t*_0_. Where warranted, we also included strain and mixing effects on *K* and *σ*. For the natural isolates, we used separate crossed factors for genetic background (A23 or A47) and antibiotic marker (Km^R^ or Rif^R^). We tested the statistical significance of model terms with likelihood ratio tests comparing the difference in deviance between nested models to a *χ*^2^ distribution.

### Mathematical model

To further test the local competition hypothesis, we constructed a mathematical model of how resource competition might occur during codevelopment, parameterized the model using time series data, and then observed how well the model predicts each strain’s sporulation success as a function of mix frequency (a separate data set).

The model describes the change over time in the number of spores *n* produced by each codeveloping strain *i*:

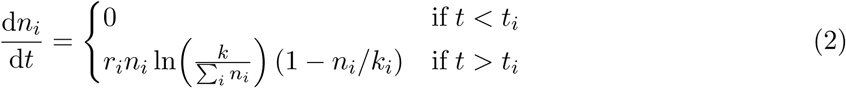

where *t*_*i*_ is the lag time until a strain begins producing detectable spores, *r*_*i*_ is a sporulation rate parameter, *k* is the sporulation carrying capacity (total group productivity), and *k*_*i*_ is the initial number of starving cells from strain *i*.

To construct the model, we first noted that Gompertz functions like Eqn. 1 are known solutions to the Gompertz model of population growth [32] and hypothesized that sporulation might follow similar dynamics. The general form of the Gompertz growth model is d*n/*d*t* = *rn* ln(*k/n*). We added a simple form of resource competition by including all strains’ spores in the denominator of the logarithm term so that net sporulation rate declines as total group spore production approaches *k*. We then included the (1 − *n*_*i*_*/k*_*i*_) term to limit a strain’s maximum sporulation success to one spore/cell. This term need not be linear, but in the absence of information this was the simplest assumption.

The value of *k*_*i*_ is determined by experimental conditions but *r*_*i*_, *t*_*i*_, and *k* must be estimated from data. With initial conditionsv ∑_*i*_ *n*_*i*_(*t* = 0) = 10 spores (the lower detection limit), *r*_*i*_ and *t*_*i*_ can be measured from developmental time series as in Eqn. 1. To get better estimates, we measured *r*_*i*_ and *t*_*i*_ using data both from strains alone and in codevelopment when statistical tests indicated these terms were unaffected by mixing. Because codevelopment with GJV10 affected GJV2’s lag time but not its sporulation success, we hypothesized that the observed differences in dynamics were an effect of rifampicin resistance on spore viability during heat treatment, sonication, and/or plating. To get better estimates of GJV2’s true sporulation dynamics we thus used *r*_GJV10_ and *t*_GJV10_. We measured *k* as the geometric mean spore production of GJV10-T developing by itself.

With these parameter values, we used the *deSolve* package in R to numerically solve Eqn. 2 for both codeveloping strains across a range of initial strain frequencies. We hypothesized that differences in total spore count between GJV10-T and GJV2 reflected spore viability and/or efficiency of resource use and thus multiplied *n*_GJV2_ by a relative viability term: (geometric mean spore count of GJV2 developing alone) / (geometric mean spore count of GJV10-T developing alone). We then calculated spores/cell for each strain at *t* = 120 hrs and compared the results to experimental data.

## Results

### Facultative social exploitation by a tan variant

We assayed social competition and exploitation using “mix experiments” that measure the sporulation success of strains when separate and when mixed together at various frequencies. To control for any effects of the antibiotic markers we used to distinguish codeveloping genotypes, we first examined interactions between two wild-type strains: kanamycin-resistant GJV10 and rifampicin-resistant GJV2 (Fig. 1A). GJV10 produced slightly more spores than GJV2 when developing separately (*t*(5.39) = 2.7, *P* = 0.039), but codevelopment did not affect either strain’s performance 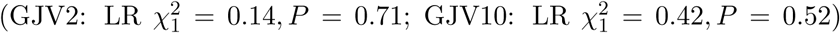 The fitness effects of the antibiotic markers were thus small and unaffected by codevelopment.

**Figure 1:**
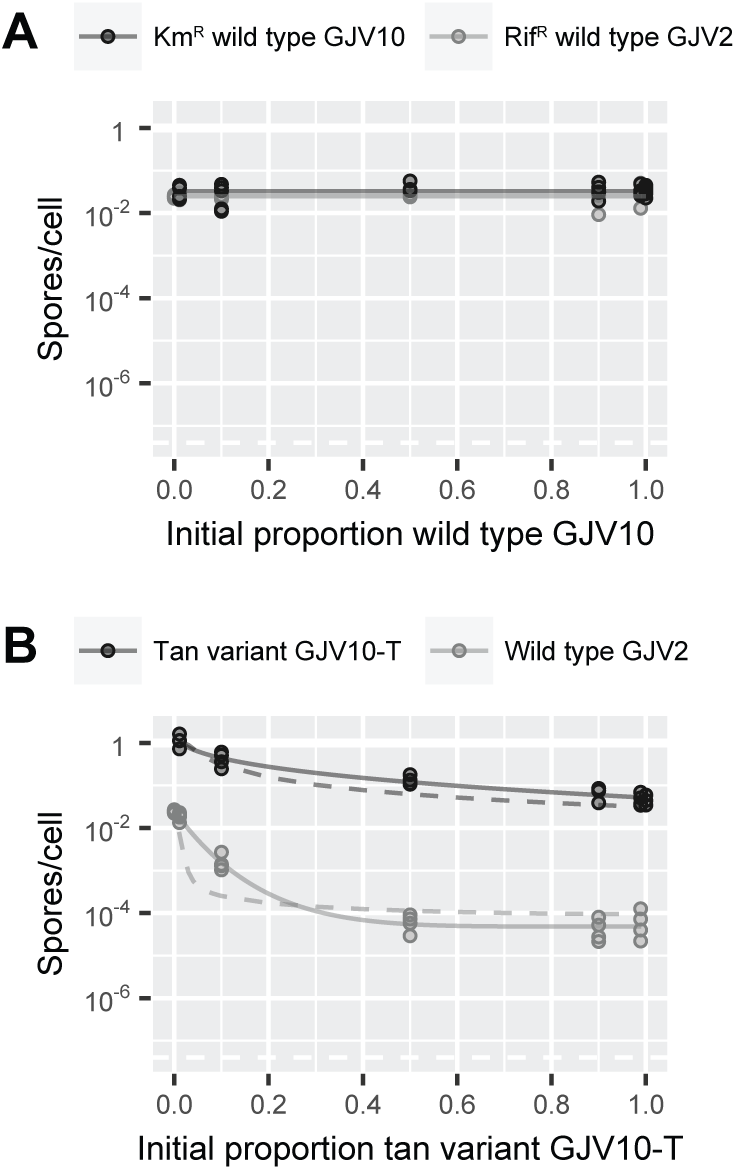
Effects of codevelopment on sporulation success of *Myxococcus xanthus* strains. Points show data from replicate experiments. Solid lines show fitted statistical models. **(A)** Marker control. Kanamycin-resistant wild type sporulates more than rifampicin-resistant, but this difference is small and unaffected by mixing. **(B)** Facultative social exploitation by tan variant. Both strains sporulate well when separate. Mixing increases absolute sporulation success of tan variant and decreases that of yellow wild-type. Dashed lines: mathematical model of local resource competition, parameterized using time dynamics of spore formation (Fig. 3), predicts magnitude and frequency dependence of mixing effect.

We isolated tan phase variant GJV10-T from wild-type yellow strain GJV10 by picking a spontaneously generated tan colony. We did not measure phase switching rates, but they were sufficiently low that revertants remained a small minority of all measured colonies for the experiments described here. The tan variant was proficient in both swarming and development, producing around twice as many more spores than its parent strain when separate (*t*(6.29) = 2.64, *P* = 0.037). In mix experiments, codevelopment with yellow wild-type GJV2 increased the absolute sporulation success of GJV10-T (Fig. 1B; *F*_2,23_ = 191.7, *P* < 0.0001) to a maximum of ∼ 1 spore/cell and decreased that of GJV2 > 100-fold 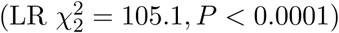. The tan phase variant thus showed facultative social exploitation, sporulating well on its own and exploiting the yellow wild-type in codevelopment.

### Group productivity unimpaired by codevelopment

To test whether social exploitation by the tan variant was due to local competition for some resource necessary to produce spores, we tested the prediction that genotype mixing would have little to no effect on total group productivity. We thus reanalyzed mix experiment data in terms of relative within-group fitness and total group sporulation success (Fig. 2). In the marker control, GJV10 had no significant within-group advantage over GJV2 (*F*_1,21_ = 2.31, *P* = 0.14). Total sporulation success of mixed-genotype groups was uniformly high, increasing slightly with GJV10 frequency 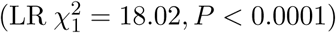. The tan variant had a large within-group fitness advantage (*F*_3,17_ = 754.9, *P* < 0.0001) that was greatest when it was common (*F*_2,17_ = 46.76, *P* < 0.0001). Total sporulation success of mixed tan/yellow groups was uniformly high, even increasing somewhat with tan abundance (*F*_2,30_ = 27.85, *P* < 0.0001). Log-sporulation success increased nonlinearly with tan frequency (*F*_1,30_ = 21.33, *P* < 0.001) but there was no significant effect past 50:50 (*F*_1,16_ = 2.61, *P* = 0.126). The tan variant therefore exploited the yellow wild type without impairing group productivity, consistent with local competition.

**Figure 2:**
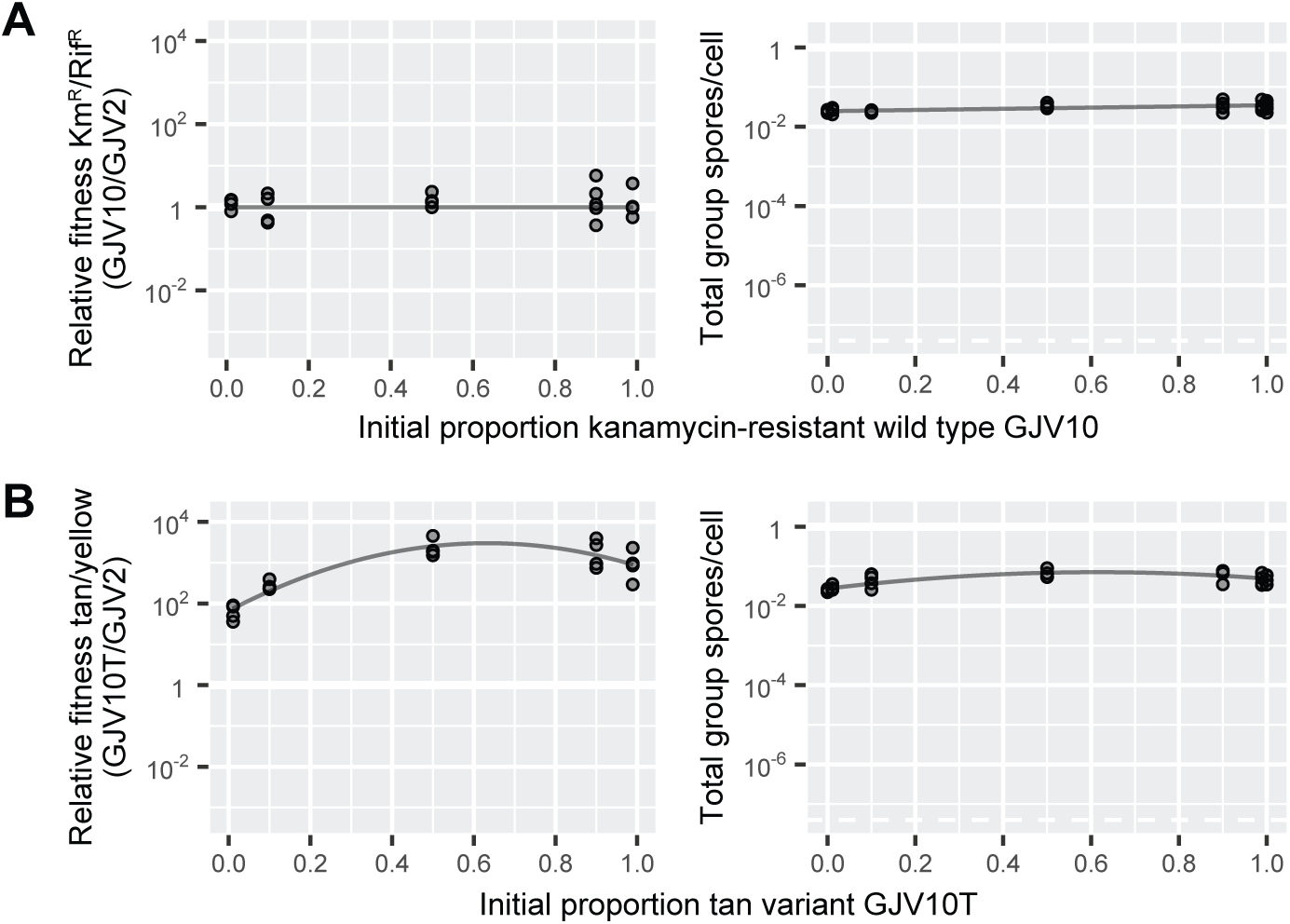
Within-group relative fitness and total group sporulation success during codevelopment. Points show data from replicate experiments. Lines show fitted statistical models. **(A)** Marker control. Little to no fitness difference between kanamycin-resistant and rifamicin-resistant wild type strains. **(B)** Tan variant outcompetes yellow wild type without impairing total group productivity.

### Time dynamics of codevelopment indicate non-responsive rate competition

We also tested whether the time dynamics of spore production were consistent with non-responsive local competition. Specifically, we tested predictions that the winning strain begins producing spores earlier and/or faster and that codevelopment does not affect sporulation rate. In the marker control (Fig. 3A), we found that GJV10 started producing spores earlier than GJV2 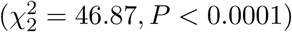 but at no faster rate 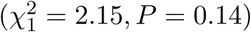. Codevelopment had no significant effect on GJV10’s lag time 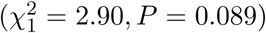 or sporulation rate 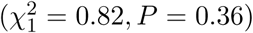.Codevelopment actually decreased GJV2’s lag time a few hours 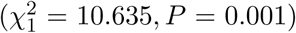 but did not affect its rate 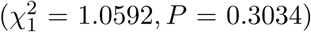. The antibiotic markers thus did have a small effect on the timing of spore production, but not in any way that created codevelopment-specific fitness effects.

**Figure 3:**
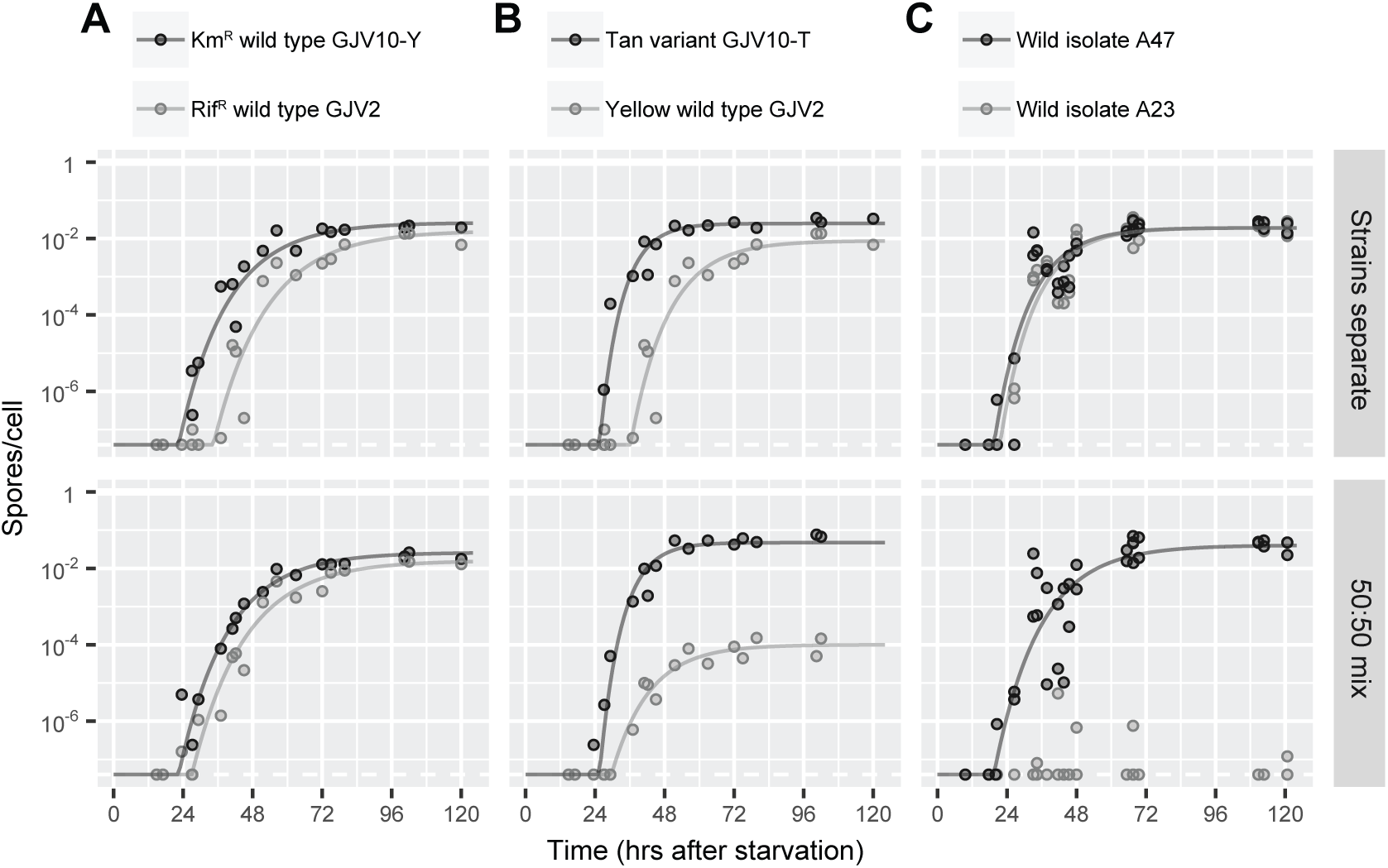
Time dynamics of developmental competition. Points show data from replicate experiments. Solid lines show fitted statistical models. Dashed white line shows lower limit of detection. **(A)** Marker control. Kanamycin-resistant wild type begins producing spores earlier and reaches a higher final level than isogenic rifampicin-resistant strain. Mixing decreases Rif^R^ time but otherwise has no effect. **(B)** Codevelopmental dynamics of tan phase variant and yellow wild type are consistent with non-responsive local resource competition. Tan sporulates earlier and faster than yellow wild type. Mixing does not affect either strain’s sporulation rate but causes yellow to stop sporulating earlier. **(C)** Dynamics of two naturally occuring soil isolates are consistent with interference competition. When separate, A47 begins producing spores slightly earlier than A23 but at the same rate. When mixed, A23 produces few to no spores and A47 sporulates at a reduced rate. Data show combined results with reciprocally marked strains (see also Fig. S3).

The tan variant showed accelerated development (Fig. 3B). GJV10-T sporulated at a faster rate than both GJV2 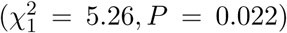 and its wild-type parent strain GJV10 (Fig. S1; 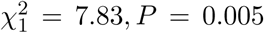). GJV10-T started producing spores earlier than GJV2 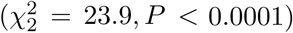, but not GJV10 (Fig. S1; 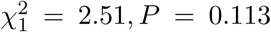). Codevelopment with GJV2 had no effect on GJV10-T’s lag time 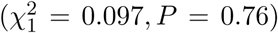 or sporulation rate 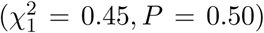.Codevelopment had no effect on GJV2’s sporulation rate 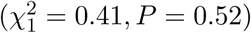, but it did decrease lag time a few hours 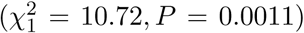, similar to the effect seen in codevelopment with wild-type GJV10. The time dynamics of tan/yellow codevelopment were therefore consistent with those predicted under non-responsive local competition.

We also examined the the codevelopmental dynamics of two natural isolates previously shown to exhibit social exploitation without a decline in total group productivity [6] through contact-dependent interference competition [19]. These strains showed facultative social exploitation in our experiments, as well (Fig. S2). When developing separately, there was no significant difference between isolates A47 and A23 in developmental rate 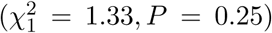 or lag time 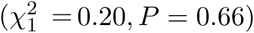, though the Rif^R^ derivative of isolate A47 did start producing spores earlier (Fig. S3; 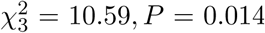). Strikingly, A23 produced almost no spores when codeveloping with A47, even in the period 24–48 hrs post starvation when total group productivity had not yet been exhausted (Fig. 3C). Codevelopment decreased the sporulation rate of A47 Km^R^ 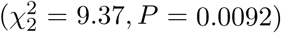 but did not affect lag time 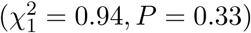. For these natural isolates, then, the time dynamics of codevelopment did not fit the predictions of non-responsive local competition and were instead consistent with interference competition.

### Mathematical model of local competition predicts frequency-dependent fitness

To further test whether social exploitation by the tan variant was due to non-responsive local competition, we constructed a mathematical model of how strains might constitutively compete for some locally shared resource necessary for sporulation, parameterized the model with time series data, and then examined how well it predicted each strain’s frequency-dependent response to mixing (a separate dataset). Given only the time at which strains begin producing spores, how fast they begin to produce spores, and the total number of spores they produce when developing by themselves, the model quantitatively predicted the codevelopmental fitness of both tan and yellow strains remarkably well (Fig. 1B). The predicted magnitude and frequency dependence of tan sporulation success closely match observed data. The model also successfully predicted the asymmetrical response of strains to codevelopment (yellow loses more than tan gains), the asymmetrical frequency dependence (steepest slopes at low tan frequencies), and the amount of spores produced by the yellow strain when the tan variant is common. The only weakness of the model was that it over-estimated how rapidly yellow sporulation success decreased with increasing tan frequency.

## Discussion

To better understand facultative social exploitation among *Myxococcus bacteria* during the formation of multicellular fruiting bodies, we tested the hypothesis that codeveloping cells compete for some locally shared resource necessary to produce spores but otherwise do not react to nor influence each other. We found strong support for this form of competition between a tan phase variant and the yellow wild-type. First, total group spore production was largely unaffected by genotype mixing, with no antagonism between strains. Second, the winner was the fastest-developing strain. Third, developmental rate was unaffected by codevelopment, with no change by either strain that did not also occur in marker-control experiments. Finally, a mathematical model of local resource competition, parameterized with time series data, successfully predicted the frequency-dependent sporulation success of both strains in codevelopment (a separate dataset). Some authors have speculated that tan *M. xanthus* cells serve some vital prespore role in development [21], but our data instead suggest that tan cells simply develop more quickly.

Our results illustrate how facultative social exploitation among microbes, in the sense of an interaction between two high-fitness strains that increases the absolute fitness of one genotype at the expense of another, does not require elaborate mechanisms to detect and respond to foreign genotypes. Instead, it can be caused by simple competition acting at a local scale. Another constitutively-expressed mechanism for facultative social exploitation occurs in *Dictyostelium* amoebae, where strains vary in both production of and sensitivity to secreted substances that induces cells to become spores [33]. Competition is being increasingly recognized as pervasive in the microbial world, even in interactions once thought cooperative [34, 35]. We note that an important role for competition does not diminish the usefulness of *Myxococcus* or *Dictyostelium* as models of microbial cooperation. In social insects, the production of reproductive castes is the outcome of both cooperative traits like foraging or brood care and competitive traits like dominance or siblicide [34].

Our data do not identify what locally shared resource tan and yellow cells compete over. The cells in our experiments were placed on non-nutrient agar and began sporulating due to starvation, so the resource seems likely to be something produced by the cells themselves. Tan phase variants upregulate genes involved in acquiring iron [23], so iron released by dying *M. xanthus* cells is one candidate. Another possibility is that there is no physical resource and local competition is instead mediated by a developmental carrying capacity. If the proportion of cells that become spores is regulated via some “first come, first served” process, strains that develop faster might be better able to find a place among the survivors. In *Dictyostelium* amoebae, which also form multicellular fruiting bodies, the first cells to begin developing preferentially become spores [35].

Our previous theoretical treatment described facultative social exploitation due to non-responsive local competition as part of a purely nonadaptive process in which high genetic relatedness relaxes selection against some kinds of deleterious mutation [17]. The situation with tan variants appears more complicated. The mutation examined here is beneficial in codevelopment, not deleterious. Tan variants are also more susceptible to ultraviolet light [36] and nematode predation [22], so these mutations appear to have antagonistic pleiotropic effects on different components of fitness. Naturally co-occuring genotypes of *M. xanthus* show similar pleiotropic effects, with some isolates performing well during vegetative growth but sporulating poorly [37]. The net effect of tan mutations could be deleterious, as most newly-formed phase variants appears to be [38]. Or it could be a bet-hedging adaptation for dealing with fitness trade-offs in unpredictable environments [39, 40, 41] where cells stochastically adopt cooperative or exploitative strategies, reminiscent of Robert Louis Stevenson’s fictional Dr. Jeckyll and Mr. Hyde [42]. Without data measuring selection on tan variants in natural populations, though, the apparent simplicity of the tan phenotype means that it does not yet meet George Williams’ standard that a biological effect should only be claimed adaptive if “it is clearly produced by design and not by chance” [15, p. v].

We also investigated the time dynamics of codevelopment between two naturally occuring soil isolates. Antagonism between these strains is caused at least in part by contact-dependent interference competition [19]. We found that the winning strain almost completely prevented the loser from producing any spores, even in the early stages of development when total group productivity had not yet been exhausted. Because the victor kills loser cells during vegetative growth [19] and developing cells feed on the lysed [43], facultative social exploitation among natural *Myxococcus* isolates may often be a simple matter of killing and eating one’s competitors. And since exploitation among laboratory-evolved strains is not associated with faster morphological development [8], there appear to be many mechanisms of social competition in *Myxococcus*.

Our work also illustrates how studying the time dynamics of competition, instead of just its beginning and end points, can provide a rich source of data for testing hypotheses about mechanisms of social selection. Population ecologists use this dynamical approach extensively and have developed many of the relevant statistical and mathematical methods [31]. Similar approaches have been used to study social exploitation in yeast [44] and antibiotic-resistant *E. coli* [45], but there is still potential for wider use. The key is developing mathematical models that have enough biological detail to make specific testable predictions.

## Acknowledgments

We thank Suegene Noh for statistical discussion.

## Data, code, and materials

The data associated with this manuscript are archived in Dryad (doi:10.5061/dryad.**TBD**).

## Author contributions

js conceived the study, designed the study, performed the experiments, analyzed the data, created the mathematical model, and drafted the manuscript. GJV provided ongoing feedback on study design, experimental results, and manuscript drafts. All authors gave final approval for publication.

## Supplemental Material

**Figure S1:**
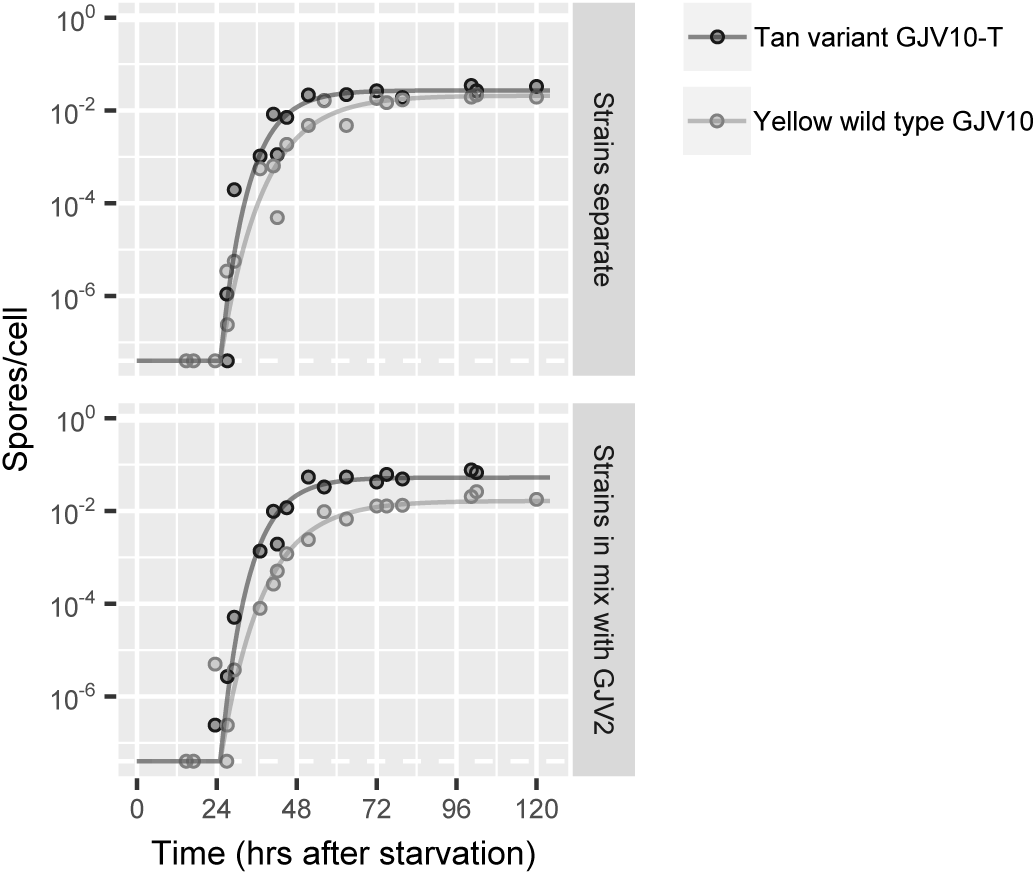
Tan variant GJV10-T sporulates faster than its yellow parent strain GJV10 but has the same lag time. Points show data from replicate experiments and solid lines show fitted statistical models. Dashed white line shows lower limit of detection.

**Figure S2:**
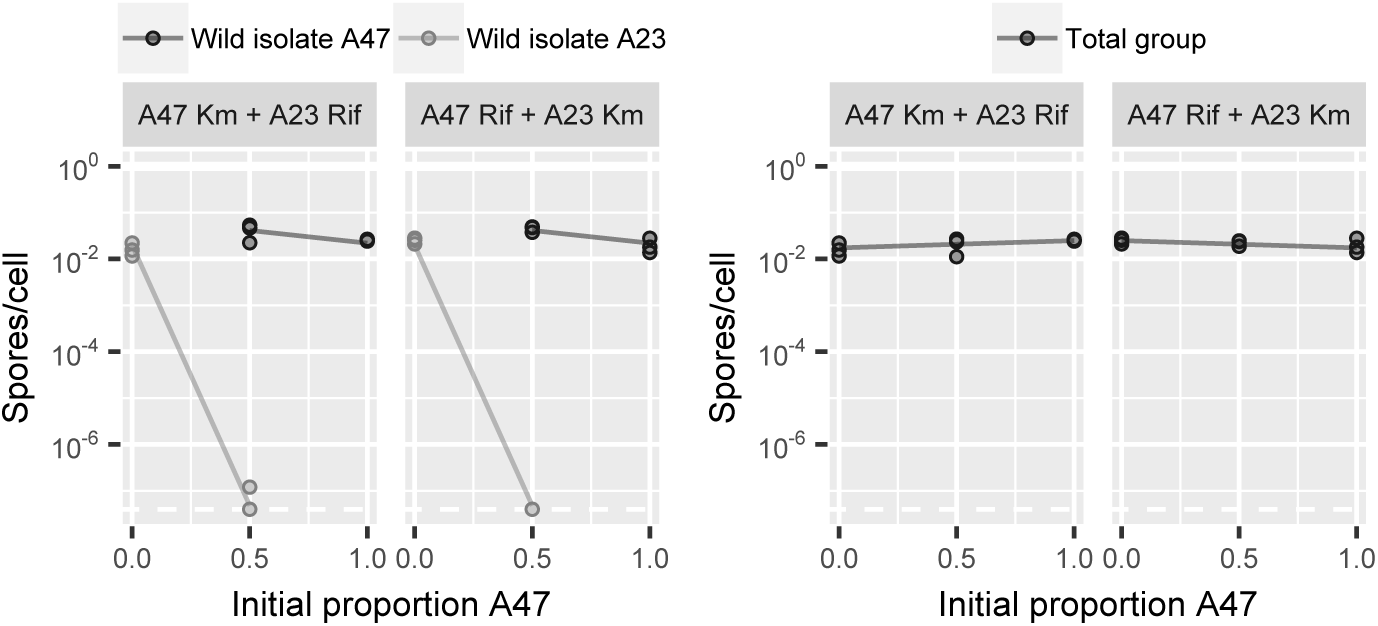
Facultative exploitation of wild isolate A23 by isolate A47. Data show last time point of sporulation time courses. Lines show fitted statistical models. Mixing increases sporulation success of A47 and decreases that of A23 without impairing total group productivity. Rifampicin resistance slightly reduces total group productivity.

**Figure S3:**
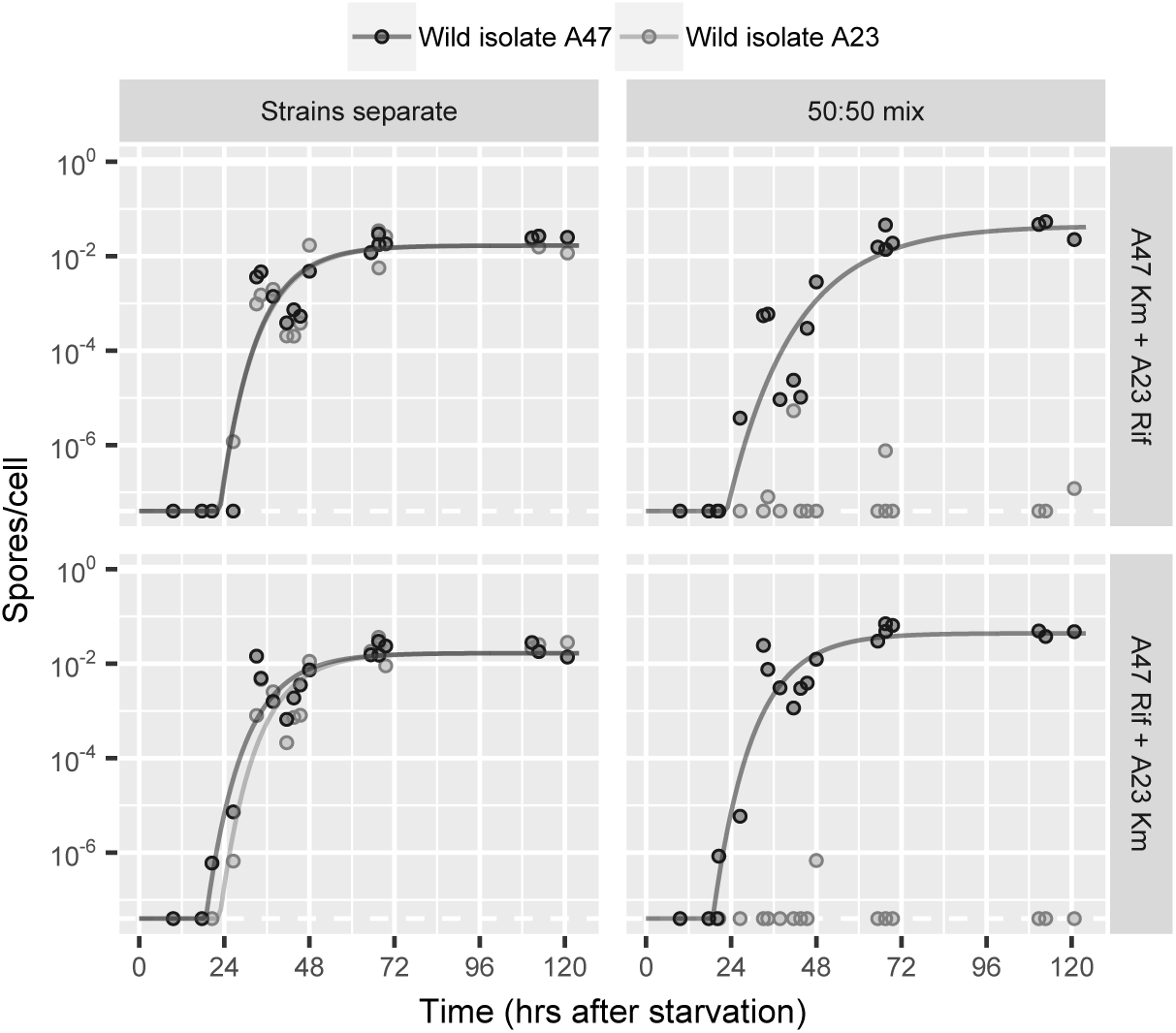
Developmental competition between natural isolates A47 and A23, with data separated by antibiotic marker. A47 Rif^R^ starts sporulating slightly earlier than A47 Km^R^. A47 Km^R^ sporulates at a reduced rate during codeveopment with A23. Data points show replicate experiments and solid lines show fitted statistical models. Dashed white line shows lower limit of detection.

